# Predicting the timing of ecological phenomena across regions using citizen science data

**DOI:** 10.1101/2023.05.05.539567

**Authors:** César Capinha, Ana Ceia-Hasse, Sergio de-Miguel, Carlos Vila-Viçosa, Miguel Porto, Ivan Jarić, Patricia Tiago, Nestor Fernandez, Jose Valdez, Ian McCallum, Henrique Miguel Pereira

**Author notes:** Corresponding author: César Capinha | @.

## Abstract

Spatial predictions of intra-annual ecological variation enhance ecological understanding and inform decision-making. Unfortunately, it is often challenging to use statistical or machine learning techniques to make such predictions, due to the scarcity of systematic, long-term observational data. Conversely, opportunistic time-stamped observation records, supported by highly informative data such as photographs, are increasingly available for diverse ecological phenomena in many regions. However, a general framework for predicting such phenomena using opportunistic data remains elusive. Here, we introduce a novel framework that leverages the concept of relative phenological niche to model observation records as a sample of temporal environmental conditions in which the represented ecological phenomenon occurs. We demonstrate its application using two distinct, management-relevant, ecological events: the emergence of the adult stage of the invasive Japanese beetle (*Popillia japonica*), and of fruiting bodies of the winter chanterelle mushroom (*Craterellus tubaeformis*). The framework accounts for spatial and temporal biases in observation data, and it contrasts the temporal environmental conditions (e.g., in temperature, precipitation, wind speed, etc.) associated with the observation of these events to those available in their occurrence locations. To discriminate between the two sets of conditions, we employ machine-learning algorithms (boosted regression trees and random forests). The proposed approach can accurately predict the temporal dynamics of ecological events across large geographical scales. Specifically, it successfully predicted the intra-annual timing of occurrence of adult Japanese beetles and of winter chanterelle mushrooms across Europe and North America. We further validate the approach by successfully predicting the timing of occurrence of adult Japanese beetles in Northern Italy, a recent hotspot of invasion in continental Europe, and the winter chanterelle mushroom in Denmark, a country with a high number of records of this mushroom. These results were also largely insensitive to temporal bias in recording effort. Our results highlight the potential of opportunistic observation data to predict the temporal variation of a wide range of ecological phenomena in near real-time. Furthermore, the conceptual and methodological framework is intuitive and easily applicable for the large number of ecologists already using machine-learning and statistical-based predictive approaches.

## Introduction

Ecological phenomena with intra-annual variation, such as species phenology, migrations, behaviour, or productivity levels, are key drivers and indicators of the structure, status and functioning of ecological systems (Tang et al., 2016). Spatial predictions of such phenomena over short and long time frames now serve a variety of important fundamental and applied purposes, including improved understanding of ecological processes (Dietze et al., 2018; Houlahan et al., 2017), anticipation of ecological risks (e.g., Kim et al., 2023), management of threats to biodiversity (e.g., Henden et al., 2022; Slingsby et al., 2023), and the promotion of sustainable use of natural resources (e.g., Marolla et al., 2021). These contributions are of growing significance given escalating environmental changes and mounting human pressures on biodiversity (Cardinale et al., 2012).

Predictions of ecological phenomena that change over time are typically derived from either process-based or data-driven modelling approaches. The first approach relies on *a priori* knowledge about the identity and type of relationship between driving factors (e.g., weather) and the temporal responses of the phenomena of interest. This knowledge is explicitly incorporated into the models using statistically derived or deterministic equations, for example by using thermal forcing units (a measure of accumulated temperature) above or below certain thresholds the phenomena is considered to occur (Taylor & White, 2020). While these models can offer high potential reliability, their implementation is dependent on the availability of in-depth prior knowledge about driving factors and ecological responses, which may be too complex or expensive to obtain for many ecological phenomena (e.g., Hassall et al., 2017). The second approach is purely statistical or machine learning-based, (i.e., so-called “data-driven”). Unlike process-based models, this approach relies entirely on algorithmic-based identification of predictive features, either in the temporal progression of the event itself or in putative environmental drivers. While this approach is generally straightforward and requires less prior ecological knowledge than the process-based approach, its use is even more dependent upon the availability of observational data amenable to model fitting. More specifically, commonly used data-driven modelling techniques, such as state-space models or custom-built machine learning architectures are mostly fitted using time series of the event, preferably collected over representative geographical extents (e.g., Lofton et al., 2022; Marolla et al., 2021; Rammer & Seidl et al., 2019; Morera et al., 2021). Unfortunately, datasets meeting these requirements are often non-existent or remain temporally or spatially limited for many ecological phenomena.

At the same time, the number of biodiversity observation records in public repositories, such as GBIF (gbif.org) or iNaturalist (inaturalist.org), has been rising steeply (GBIF, 2023). These data, particularly those from citizen-science platforms, are frequently georeferenced with high precision, time-stamped, and accompanied by visual media, including photographs. As such, they represent a potentially valuable source of information on the spatiotemporal dynamics of ecological phenomena. Previous research has already demonstrated their usefulness for temporal ecology research, such as in measuring flight periods for Lepidoptera species (Belitz et al., 2023) or in estimating the flowering period of plant species (Puchałka et al., 2022). However, despite their widespread availability, the use of presence-only, predominantly opportunistic data for temporal modelling is challenging due to the lack of temporal replicability, temporal recording biases, and uneven spatial coverage. To minimise these limitations, previous works have selected records from areas with temporal replicability (e.g., where multiple records are available within the same year) from which temporal trends are then interpolated (e.g., Belitz et al., 2020; Puchałka, et al., 2022). Although practical and seemingly effective (Belitz et al., 2020; Pearse et al., 2017), this approach disregards potentially informative records in regions with low or null temporal replicability. Moreover, the need for temporal replicability also creates data availability issues, similar to those of time series data, limiting the phenomena and regions that can be modelled.

In this study, we propose a novel, data-driven, approach for predicting the intra-annual timing of occurrence of ecological phenomena using opportunistic presence-only records. The approach is grounded in ecological theory and assumes that any observation record of the phenomenon of interest reflects the temporal match of physical and biological conditions suitable for its occurrence. By jointly sampling the set of conditions represented across multiple records, our approach constructs a representation of the temporal environmental space under which the phenomenon occurs, i.e., its ‘phenological niche’ (Post, 2019). This approach can integrate all occurrence data available and is not reliant on regional temporal replicability. To demonstrate its effectiveness, we use it to provide daily predictions of the occurrence of adult invasive Japanese beetles (*Popillia japonica*) and fruiting bodies of the winter chanterelle mushroom (*Craterellus tubaeformis*) across Europe and North America. We also show its applicability for management-related tasks by using environmental predictors enabling the near-real-time prediction of these two ecological events. Our approach is conceptually intuitive and straightforward to implement for ecologists experienced with machine learning or statistical-based predictive modelling. It also provides a promising research avenue to harness the vast and growing amounts of presence-only opportunistic data for predicting the timing of ecological phenomena.

## Materials and Methods

### Conceptual framework

In this section, we introduce the theoretical basis for the components represented in our modelling framework and the estimates that the models provide. While our framework can be applied to a range of temporally varying ecological phenomena, we base our conceptual framework on the concept of the ’phenological niche’ (Post, 2019), as much of the core theory of temporal modelling in ecology originates from the field of phenology. Post (2019) differentiated between the ’absolute’ and ’relative’ phenological niche. The former refers to the use that species make of absolute or ’cosmological’ time, such as distinguishing between ’early’ or ’late’ life-history strategists. The latter, which is the focus of our framework, refers to the timing of phenological events as a function of temporal variation in biotic and abiotic factors, i.e., ’relative’ to environmental drivers.

Under this framework, relative phenological niches comprise the set of temporal environmental conditions within which a phenological event occurs, a concept that can be represented as an *n*-dimensional hypervolume (Figure 1). For example, if a hedgehog is active on a specific day of the year, this means that the conditions for relevant biotic (e.g., food-resources) and abiotic (e.g., temperature and precipitation) factors – with reference to the specific day and place of the observation – are within the boundaries of the hypervolume of ‘hedgehog activity’. Relevantly, these hypervolumes can be empirically sampled using records of the observation of the event, where each record represents a point in its *n*-dimensional space. The comprehensiveness of the sampling inherently depends on the representativeness of the set of observation records. However, with many phenomena now being represented by thousands or tens of thousands of these records (Klinger et al., 2023) it seems plausible to assume that a good representativeness can often be achieved.

**Figure 1.**
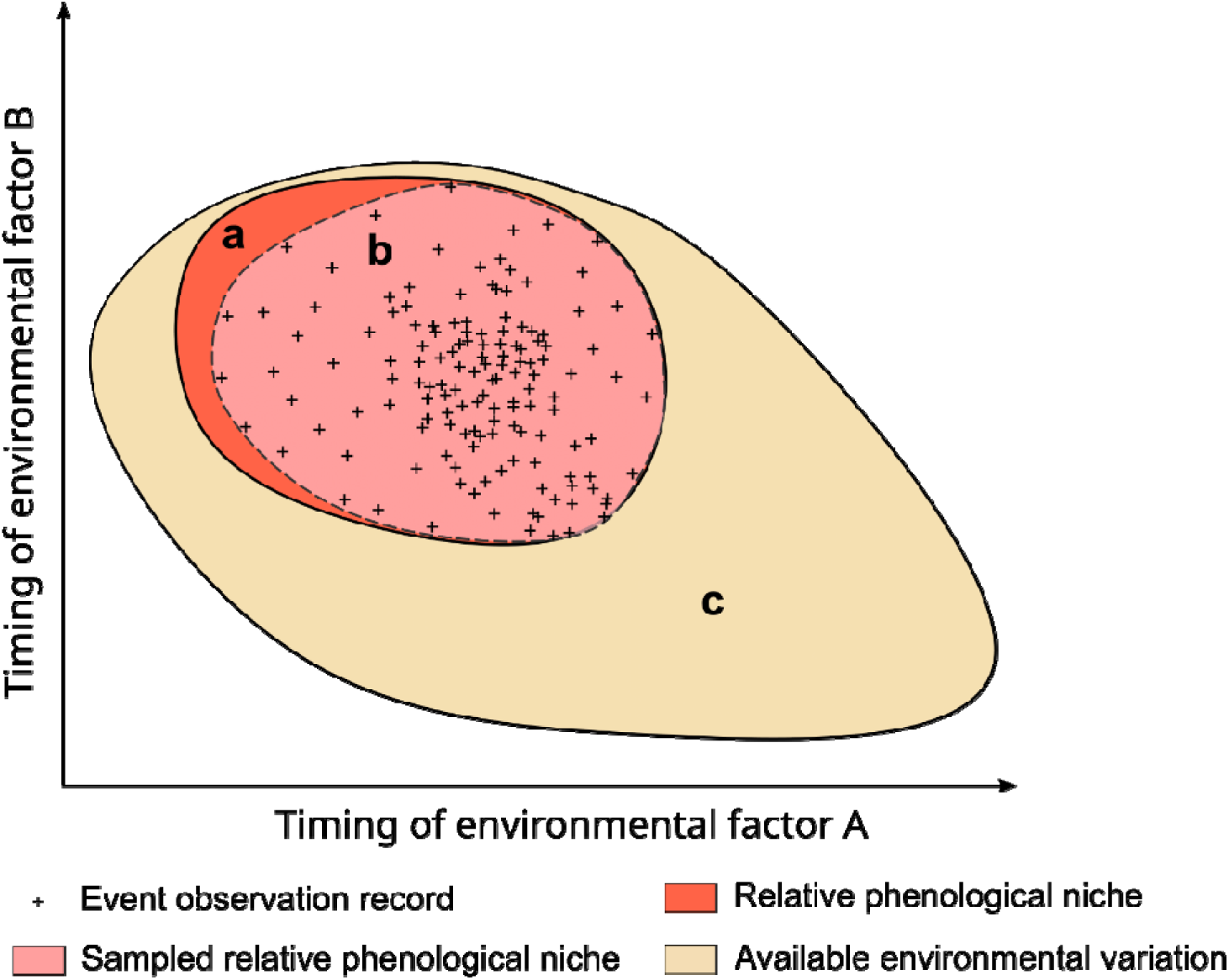
Schematic representation of the conceptual framework underlying the modelling approach. Hypervolumes of temporal environmental conditions are represented describing the relationship between the relative phenological niche of a hypothetical phenomenon of interest (a), a subset of this niche that is represented by available presence-only observation records (b), and the full set of temporal environmental variation that is available in locations where the phenomenon occurs (c). Delineation of hypervolumes is made along a simplified two-dimensional space defined by the timing of two hypothetical environmental drivers.

Empirical evidence has consistently shown that ‘group-discrimination’ techniques (e.g., random forests or regression models), which contrast the representation of one class with representations of other classes, generally perform better than alternatives which rely only on records of interest (e.g., Morera et al., 2021; Valavi et al., 2022). Hence, to allow using this class of models, we also consider the assembly of a set of environmental conditions to contrast with those representing the relative phenological niche. For that purpose, we use the set of temporal environmental conditions that are available in places where the phenomenon occurs, i.e., the so-called ‘realized environmental conditions’ (Post, 2019; Figure 1). These conditions can be sampled using records with the same geographical coordinates as observation records (guaranteeing that the sampling is made in areas where the event occurs) but with dates drawn randomly. Hereafter, we call these records ‘temporal pseudo-absences’ as they are conceptually similar to ‘pseudo-absence’ records used in species distribution modelling for sampling the geographical space available (Phillips et al., 2009).

In summary, our framework assembles a dataset representing the temporal environmental conditions associated with observation of the phenomenon of interest and with the full set of conditions in places where the phenomenon occurs. Discriminative algorithms are then used to distinguish between these two sets, and predictions can be interpreted as the probability of the represented conditions belonging to the relative phenological niche of the represented phenomenon.

### Steps of the methodological framework

In order to provide a clearer understanding of the methodology described below, we outline the main steps of our framework as follows: 1) compilation of a data set of georeferenced and timestamped occurrence observations; 2) gathering of spatial time-series of environmental variables as predictors; 3) implementation of procedures to minimize effects of spatial and temporal recording bias (optional); 4) extraction of environmental values at each location and date for the observations and temporal pseudo-absences; 5) use the resulting data set to train modelling algorithms distinguishing between environmental conditions associated with occurrences and temporal pseudo-absences; 6) evaluate the predictive performance of the models; and 7) use models to predict the occurrence of the event in places and time periods of interest. We provide a detailed description of each step below.

### Demonstration of the methodological approach using empirical data

To demonstrate the implementation of the methodological framework, we use phenological events related to the Japanese beetle and the winter chanterelle mushroom. The Japanese beetle is a highly problematic invasive species known to feed on hundreds of plant species, causing significant economic losses (Potter et al., 2002). This beetle is well established in North America and the Azores and has recently established itself in northern Italy, raising concerns of a rapid invasion of Europe from there (EFSA et al., 2019).

Early detection surveys can be effective in preventing the spread of this species, and the adult life stage is particularly suitable for such surveys (EFSA et al., 2019). Therefore, it is essential to understand when adults of this species are likely to be observed, especially in areas where their presence is uncertain, to determine the appropriate timing for implementing surveillance efforts.

The winter chanterelle is a popular edible mushroom found in Europe and North America. In some areas of these regions, the harvesting of wild edible mushrooms is regulated to avoid excessive human pressure on areas of occurrence (Copena et al., 2022). Importantly, the timing and abundance of mushroom fruiting bodies determine the level of human pressures (Górriz-Mifsud et al., 2017), thus having prior knowledge about the timing of their occurrence can aid in management decisions.

### Step 1: Assembly of event observation data

We obtained observation records for both species from the Global Biodiversity Information Facility (GBIF; https://www.gbif.org/), which is a leading aggregator of biodiversity observation records, including those from citizen science platforms such as iNaturalist (https://www.inaturalist.org/). For the winter chanterelle, we also included records from Mushroom Observer (https://mushroomobserver.org/), another citizen science platform not included in GBIF. We limited our dataset to records with photographic evidence, full date of observation (i.e., day, month, and year), and geographic coordinates with a spatial precision greater than 0.1 decimal degrees (∼11 km at the equator). We included records from 2015 to 2021 and, to ensure data quality, we removed GBIF records where the observation date was the first day of the month and the observation time was ’00:00:00’. These records generally only provide the month and year and are assigned the first day of the month by default (Belitz et al., 2023). We then assessed the photographic evidence supporting each remaining record. For the Japanese beetle, we only retained records where the photograph showed an adult life stage and no signs of the specimen being dead. For the winter chanterelle, we kept records supported by photographs of fruiting bodies that showed no signs of significant deterioration.

### Step 2: Environmental data

The timing of ecological events can be influenced by a multitude of biotic and abiotic factors. However, for the two phenomena being modelled, weather-related factors are believed to be the main drivers of their seasonality, as evidenced by previous studies (Diez et al., 2013; EFSA et al., 2019). Therefore, we used daily spatial time series of minimum temperature, mean temperature, maximum temperature, total precipitation, snow depth, and wind speed to capture the environmental conditions associated with the occurrence of these events.

We sourced these data from the AgERA5 dataset (Boogaard et al., 2020), which provides daily weather maps at a spatial resolution of 0.1° (approximately 11 km at the equator). The data was collected for the period from 2014 to 2021, but its availability has a delay of approximately one month after the last day represented. Hence, to exemplify the implementation of the framework for near-real time prediction, we collected the same set of weather variables from the Global Forecast System (GFS), a weather forecast model from the National Centers for Environmental Prediction (https://www.ncei.noaa.gov). The GFS provides forecasts of weather conditions at intervals of up to 3 hours and runs four times a day at 00:00, 06:00, 12:00, and 18:00 UTC. To ensure consistency between the two data sources, we extracted the forecasted conditions for the first six hours of each model run and aggregated them to a daily resolution. We also resampled AgERA5 data to 0.25° cell size (c. 28 km at the equator), the spatial resolution of GFS data. The processing of spatial weather data was performed in R (R Core Team) using functions provided by the ‘raster’ and ‘terra’ packages (Hijmans et al., 2023).

### Step 3: Addressing spatial and temporal recording bias (optional)

Biodiversity observation data are often geographically and temporally biased, particularly opportunistically collected records (Isaac and Pocock, 2015). To minimize these biases in our models, we applied a set of procedures described next. We note, however, that this step is optional within our framework and may be omitted if there are reasons to expect that the data is not significantly impacted by recording biases.

To avoid having regions with disproportionately high numbers of records, which could potentially dominate the overall patterns in the data (i.e., geographic overrepresentation), we first randomly selected only one record per each combination of day and 0.25 × 0.25-degree grid cell, which is the resolution of our environmental data. We also accounted for overrepresentation at the regional scale by creating a regular grid of 250 × 250 km squares covering the entire study area and counting the number of records in each square. We identified squares that exceeded the upper outlier threshold (> Q3 + 1.5 × IQR, where Q3 is the upper 25% quantile, and IQR = Q3 - lower 25% quantile) and randomly selected a number of observations equal to the threshold value (i.e., Q3 + 1.5 × IQR) to address overrepresentation.

To address temporal biases in our data, we employed a benchmark taxonomic group, *Pinus* spp. (i.e., pines), which we expect to experience variability in record availability mainly due to variation in observation effort rather than changes in the taxa phenology itself. Pines display limited visual changes throughout the year, with their evergreen foliage and inconspicuous flowering and fruiting, making them a suitable choice for our benchmark group. Thus, we can assume that the temporal variation in the number of observation records for this taxon is mainly driven by extrinsic factors, such as weather conditions, day of the week, and time of the year, that affect biodiversity recording activity (Di Cecco et al., 2021).

We downloaded all observation records of *Pinus* spp. from GBIF made between 2015 and 2021, which had the full date of observation (i.e., day, month, and year) and geographic coordinates with a spatial precision greater than 0.1 decimal degrees (∼11km at the equator). As for the Japanese beetle and the winter chanterelle, we excluded records indicating the first day of the month and an observation time equal to ’00:00:00’. Additionally, we accounted for multiple records resulting from a single recording session of the same specimen by considering all records having the same date within a 100 m radius from each other as a single record. We also identified areas of 250 × 250 km that were upper outliers in record numbers and downsampled the records in these regions by randomly selecting a number equal to the outlier threshold. In total, 303,907 *Pinus* spp. observation records were retained (Fig. S1).

Next, we generated an equal number of *Pinus* spp. observation having the same geographical coordinates, but with dates generated at random within the years represented in the observation data. This allowed us to obtain a distribution of records that would be expected if observations were made randomly over time. For both types of records (i.e., observations and randomly generated dates), we then extracted the day of the week, month, average temperature of the day, total precipitation of the day, and average wind speed of the day.

We used a generalised linear model (GLM) with a binomial error distribution, to relate the two classes of records to the calendar and weather predictors. The fitted model returned sensible results, supporting an adequate capturing of temporal recording bias (see Results section).

We then applied the model to predict the level of sampling effort for the conditions associated with each record of the Japanese beetle and winter chanterelle. The values predicted represent the propensity for having more records simply because conditions are more favourable to observers (i.e., preferred days of the week, months, and weather conditions). We then accounted for this bias in the data sets of the Japanese beetle and the winter chanterelle using inverse probability weighting (Mansournia et al., 2016). Specifically, we built a second data set of observation records for each event, where the probability of each original observation being included was inversely proportional to the level of observation effort predicted. This corresponded to randomly selecting with replacement, the same number of total records, where the probability of each record being selected was defined as 1 minus the probability predicted by the model. In other words, records made under conditions less favourable for observers had greater chances of being selected, and vice versa. While we expect this procedure to minimize temporal biases in the data, we also acknowledge that it still has limitations such as, for instance, the omission of additional drivers of observation effort (e.g., national holidays). Bearing this in mind, all subsequent analyses were carried out using both the temporally ‘corrected’ observation data and the data without this correction.

### Step 4: Extract values of predictors for event records and temporal pseudo-absences

To represent the environmental conditions at the time of each observation record, we calculated a comprehensive set of 67 features (listed in Table S1 in the Supporting Information). These features represent the geographical coordinates of each record of the event and year-long to sub-weekly conditions in mean, maximum, and minimum temperature, accumulated precipitation, wind speed, and snow depth. Importantly, each feature value was calculated in reference to the date of the record, meaning that they capture environmental conditions observed in the preceding periods, e.g., days, weeks, months, year (Table S1).

As mentioned in the conceptual framework section (above), assembling a set of environmental conditions to contrast with those representing relative phenological niches allows using discriminative modelling approaches, which tend to achieve higher predictive performance. To this end, we generated a set of temporal pseudo-absences by generating, for each observation record, a set of 12 records having the same geographical coordinates but dates drawn at random from the temporal range of the observation data. The use of 12 temporal pseudo-absences per event record was determined empirically based on preliminary tests evaluating the time taken for model training and internal cross-validation values. Although this ratio allowed us to achieve good overall predictions (see Results), we acknowledge that future work could investigate this further and additional optimization may be possible. For each temporal pseudo-absence record, we extracted the same set of 67 features used to characterize event records, providing a representation of the environmental conditions available over time in locations where the species occur (Fig. 1).

### Step 5: Model training

To differentiate between the conditions associated with the timing of observation of events and the full range of conditions available, we employed Random Forests (RF) and Boosted Regression Trees (BRT), two well-performing machine learning algorithms commonly used in ecological research (Cutler et al., 2007; Elith et al., 2008). For RF, we used the randomForest function of the R package with the same name (Liaw and Wiener, 2002), specifying a total of 2000 individual trees and remaining parameters set at default values (but see below for an exception regarding ‘sampsize’). For BRT, we used the gbm.step function of the ’dismo’ package (Hijmans et al., 2017), setting a tree complexity of 3, a learning rate of 0.005, 4 internal cross-validation folds, a bag fraction of 66%, and a maximum of 7500 individual trees.

Given the high class imbalance in our datasets (i.e., 12 temporal pseudo-absences per event record), we took steps to prevent model fitting problems such as overclassification of the majority class (Valavi et al., 2022). For RF training, we used the event observation records and an equal number of randomly selected temporal pseudo-absences for fitting each tree, as allowed by the ‘sampsize’ parameter. In BRT, we assigned a relative weight of 1/12 (i.e., 0.83%) to each temporal pseudo-absence, as allowed by the ‘site.weights’ parameter. Before each model training event, we also measured the Pearson correlation coefficient among environmental variables, retaining only the minimum set of predictors with an absolute correlation value lower than 0.8 (Valavi et al., 2022) using the ‘findCorrelation’ function from the ‘caret’ package (Kuhn et al., 2008).

### Step 6: Evaluation and validation of predictive performances

To evaluate the predictive performance of models, we first measured their capacity to correctly classify event observation records and pseudo-absences. This was evaluated for each year independently, where model training used data for the remaining years (i.e., temporally independent data). We used the area under the receiver operating characteristic curve (AUC) to measure the agreement between predictions and the actual record (i.e., observation, or temporal pseudo-absence). In the context of this work, the AUC measures the probability that observation events receive higher probability values than records generated randomly over time. AUC values range from 0 to 1, where a score of 0.7 or above is considered an acceptable level of discrimination (e.g., Valavi et al., 2022).

While the above-described evaluation procedures assess the performance of each model, they do not allow for a comparison between models using temporally corrected observation data and models using uncorrected data. To allow such comparisons, we performed a second set of evaluations comparing predictions with ‘raw’ observation data (i.e., without spatial or temporal correction) for regions left out of model training and where higher data reliability can be expected. Specifically, for the Japanese beetle, we performed this validation using observation data from northern Italy (*n* = 214), a region of high relevance for invasion surveillance of the species in Europe and where the species has received significant attention in recent years (EFSA et al., 2019). For the winter chanterelle, we used data from Denmark, the country with the highest number of observation records (*n* = 460) and where the species is foraged and marketed (Gry and Andersson, 2014).

To perform this assessment, we predicted the average probability of observing the event across the study area on each day, for years with 10 or more observation records in validation regions (i.e., 2019-2021 for the Japanese beetle and 2017-2021 for the winter chanterelle). To measure the association between the timing of observation of events and model predictions, we calculated the Point-Biserial correlation between the average predicted probability in the region and the presence (coded as 1) or absence (coded as 0) of event observations using 10-day time steps. For the two evaluation procedures (i.e., temporally corrected vs uncorrected data), we measured the performance of BRT, RF, and of an ensemble model, which is the average of the predictions of the two former algorithms.

### Step 7: Mapped predictions

To showcase the potential of our framework and assess the spatial patterns of temporal change in our predictions, we generated daily prediction maps for both events (Japanese beetle and winter chanterelle) in Europe for the year 2021. Our predictions were produced using the ensemble model trained with spatially and temporally corrected observations and AgERA5 predictor data. To avoid extrapolating beyond sampled environmental conditions, we masked all regions deemed unsuitable for the Japanese beetle (cf. EFSA et al., 2019) and regions outside the distribution range of the winter chanterelle, as represented by its observation data. To enhance visualization, we used bilinear interpolation to downsample predictions to a resolution of 0.02° (∼2km) using the ’resample’ function provided by the ’raster’ package.

## Results

### Data

In total, we obtained 15,529 event observation records for the Japanese beetle and 3,057 for the winter chanterelle, of which 10,308 and 1,726 were kept for model training (respectively) after accounting for geographic overrepresentation (Figs. S2 and S3). For both events, the number of records increased substantially along the years, with 2020 and 2021 holding more records than the previous 5 years (2015 to 2019) combined (Fig. S1a). Records of the Japanese beetle were distributed in regions of Asia, Europe, Central America, and North America, but with the vast majority concentrated in the latter, more specifically in the USA (Fig. S2 c-d). For the winter chanterelle, records were almost entirely distributed in Europe and North America, except for a few records in Central America and Japan (Fig. S3 b-d).

### Temporal bias correction

The GLM model used to examine the relationships between the availability of records for the benchmark taxa (*Pinus* spp.) with calendar- and weather-related variables yielded convincing results. The model revealed a significant (α = 0.05) positive relationship between record availability and warmer days, low precipitation, and low wind intensity (Table S2). It also identified a significantly higher propensity for observation records to be made during the weekend, on Wednesdays, and in May, June, July, and August, in comparison to Friday and April (day of the week and month used as reference level, respectively).

Conversely, significantly lower numbers of records were identified for all remaining months (January, February, March, September, October, November, and December), as well as for Mondays and Tuesdays.

### Predictive performances and mapped predictions

The BRT and RF algorithms, along with the ensemble model, consistently demonstrated excellent predictive performance when evaluated on years that were not included in the model training, achieving an area under the curve (AUC) of 0.81 or higher (Tables 1 and 2). The models for the Japanese beetle exhibited a higher discrimination capacity (average AUC = 0.9 S.D. ±0.1) than those for the winter chanterelle (average AUC = 0.85 S.D. ±0.2). Models trained on temporally corrected and uncorrected data demonstrated similar levels of accuracy overall. However, there was an apparent trend for the latter to provide slightly better results for more recent years, while the opposite was observed for earlier years. The residual values displayed significant variation across the study areas (Figs. S4 to S7), indicating that classification errors were not spatially clustered, and the global AUC values were geographically representative.

**Table 1.**
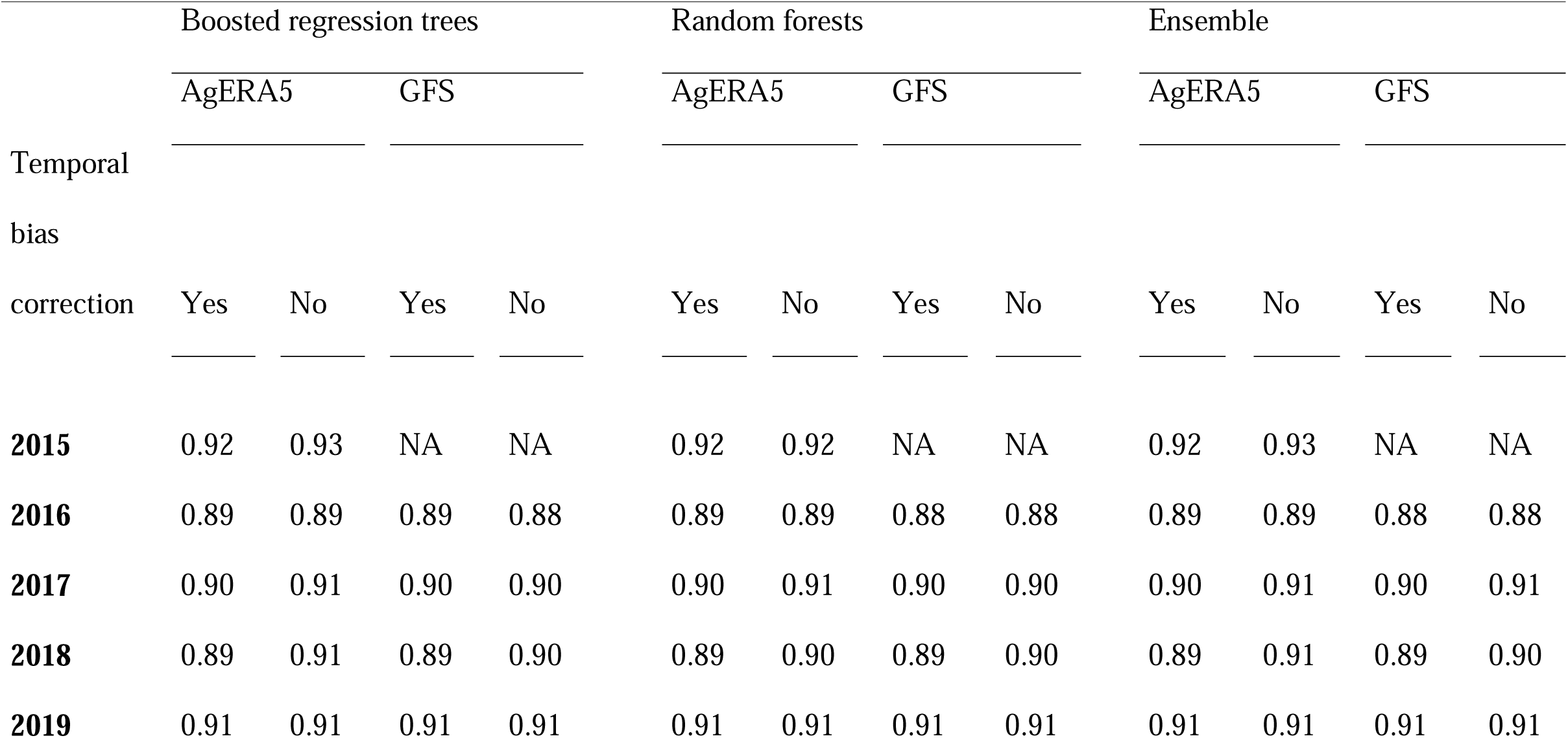
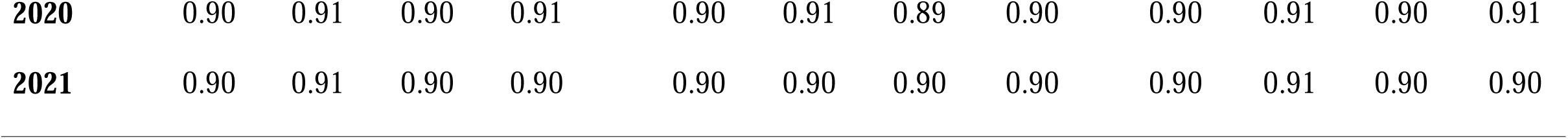
Values of area under the curve (AUC) for models predicting the timing of occurrence of adult Japanese beetles (*Popillia japonica*) across the whole study area, from 2015 to 2021. Predictions are compared with data for years that were excluded from model training. The AUC values are shown for models trained with observation data corrected for temporal and spatial bias, and for spatial bias only, for models using AgERA5 weather data and Global Forecast System data (GFS).

**Table 2.**
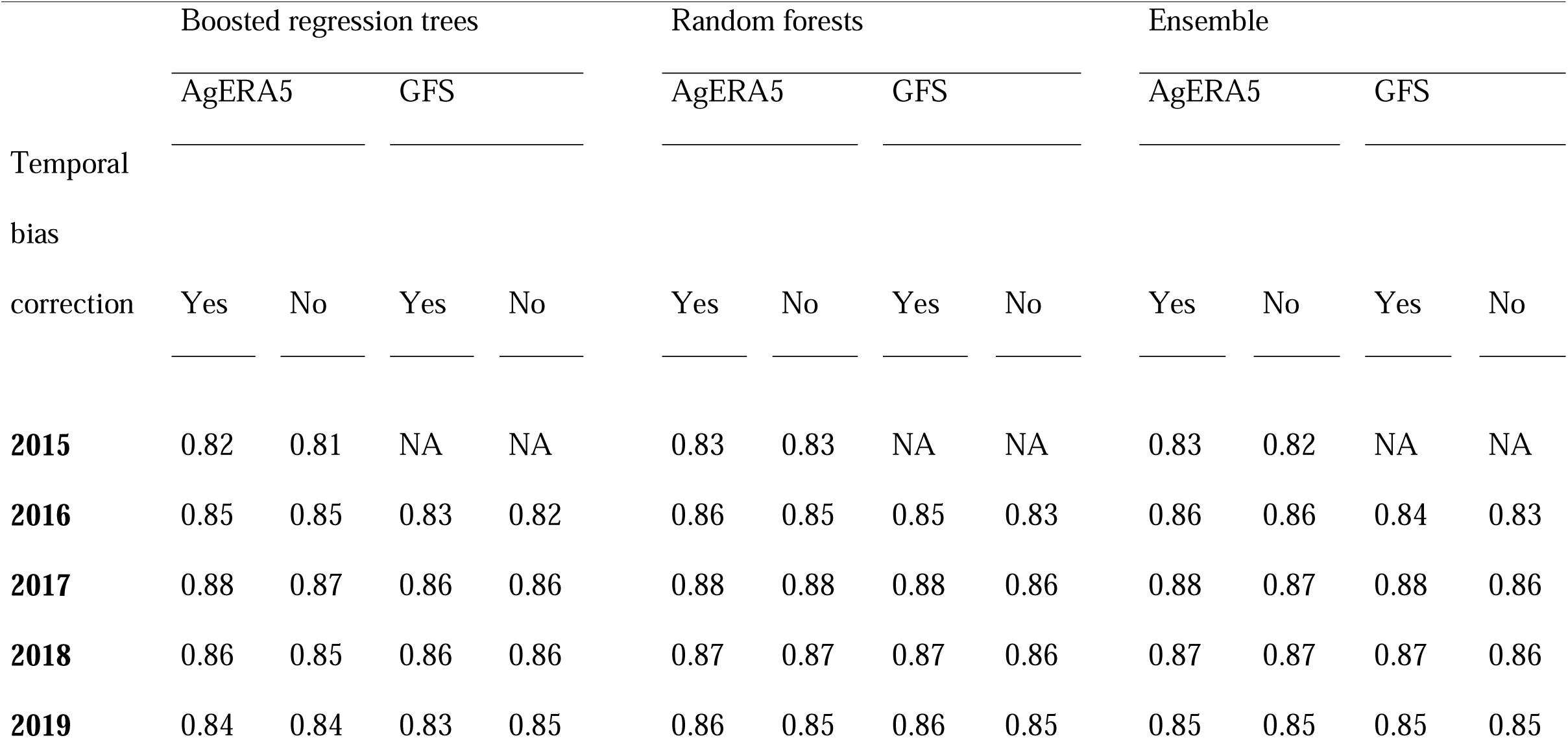
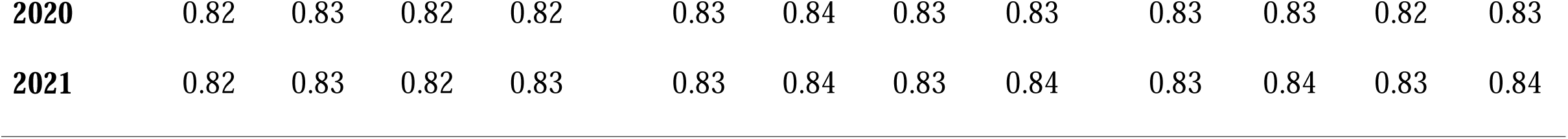
Values of area under the curve (AUC) for models predicting the timing of occurrence of fruiting bodies of the winter chanterelle (*Craterellus tubaeformis*) across the whole study area, from 2015 to 2021. Predictions are compared with data for years that were excluded from model training. The AUC values are shown for models trained with observation data corrected for temporal and spatial bias, and for spatial bias only, for models using AgERA5 weather data and Global Forecast System data (GFS).

Model evaluation in selected regions also demonstrated high predictive performance and sensible predictions. For the Japanese beetle in northern Italy, the predicted values exhibited a strong correlation with the timing of observations, with a correlation coefficient of 0.8 or higher (Fig. 2). Despite slight inter-annual variations, the results consistently indicated higher probabilities of adult beetles in mid-July (c. the 200th Julian day) and a period of higher probability of approximately 2.5 to 3 months centred on this date. For the winter chanterelle in Denmark, the correlations between predictions and observations were lower but remained very strong with *r*-values of 0.67 or higher (Fig. 3). Across years, the predicted values showed a decreasing trend from early days until the beginning of August, after which they increased rapidly, reaching a peak phase that lasted approximately between September to December (c. Julian days 244 to 365).

**Figure 2.**
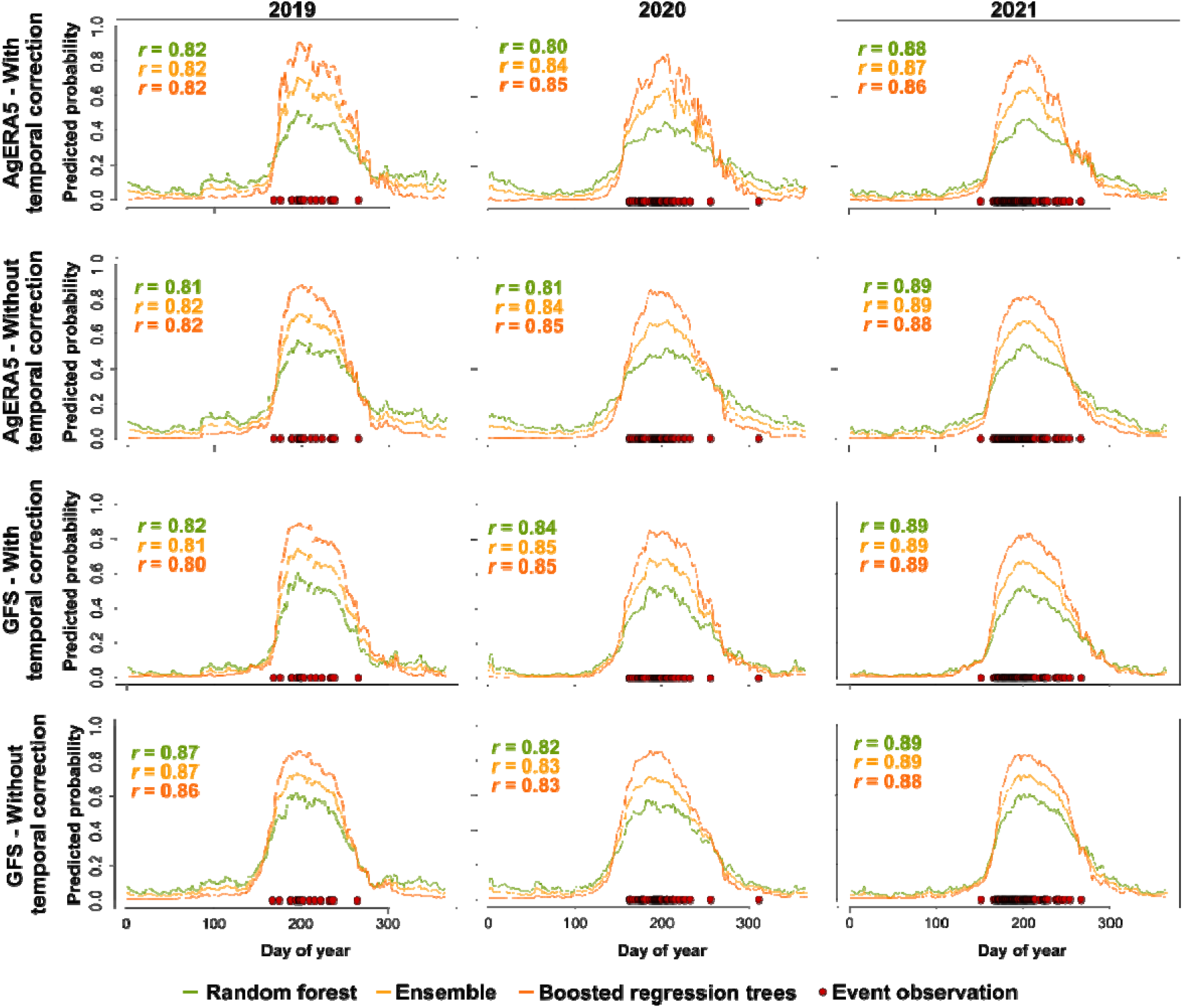
Continuous predictions of the timing of occurrence of adult Japanese beetles (*Popillia japonica*) in northwest Italy for years between 2019 to 2021. Predictions are shown for models trained with observation data corrected for temporal and spatial bias, and for spatial bias only, for models using AgERA5 weather data and Global Forecast System data (GFS). Values of Point Biserial Correlation coefficient (*r*) are provided, measuring the association between predicted values and the dates of actual observations.

**Figure 3.**
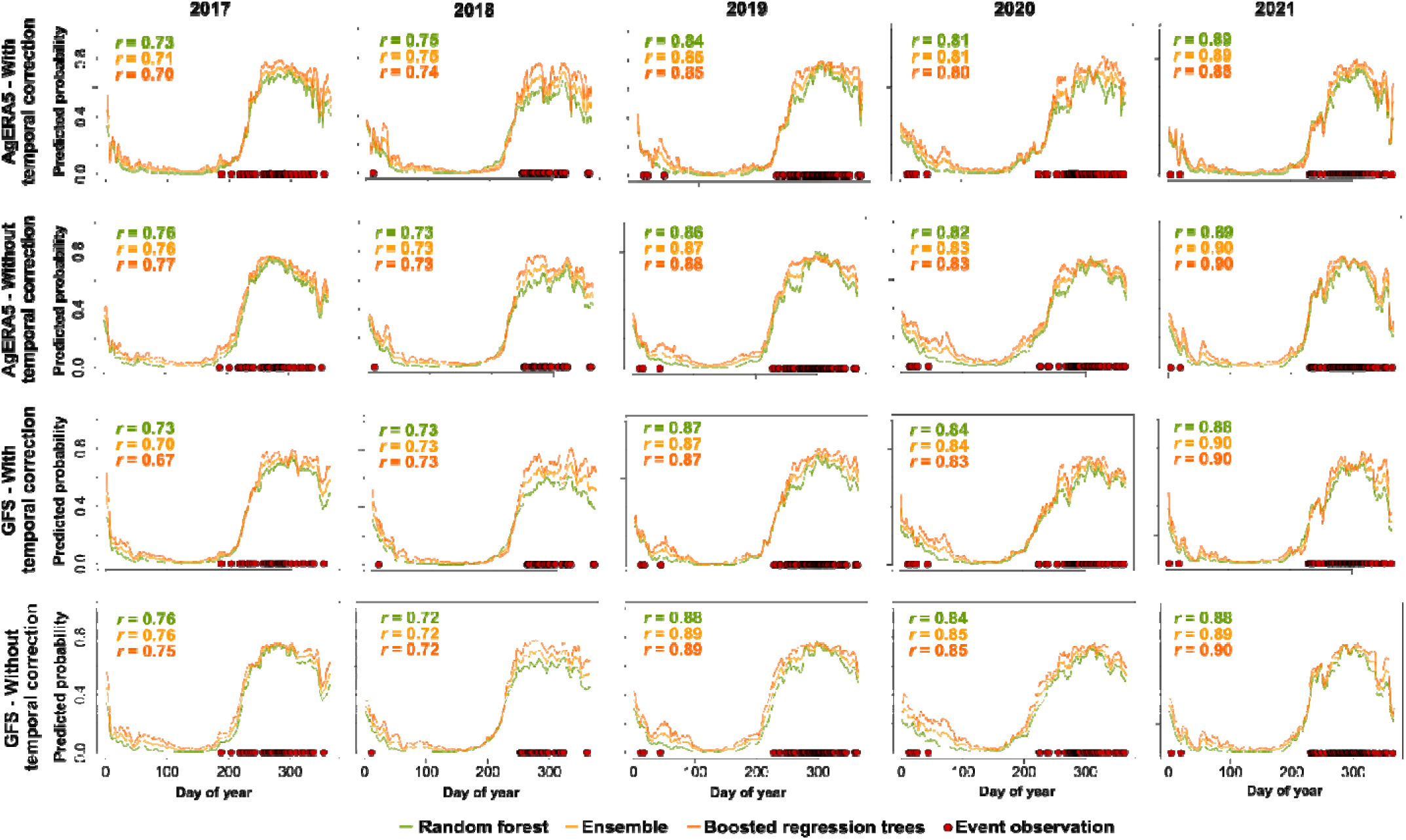
Continuous predictions of the timing of occurrence of fruiting bodies of the winter chanterelle (*Craterellus tubaeformis*) in Denmark from 2017 to 2021. Predictions are shown for models trained with observation data corrected for temporal and spatial bias, and for spatial bias only, for models using AgERA5 weather data and Global Forecast System data (GFS). Values of Point Biserial Correlation coefficient (*r*) are provided, measuring the association between predicted values and the dates of actual observations.

Maps of predictions for the Japanese beetle across Europe in 2021 show that adult beetles emerge earlier in southern regions (southern Iberia, southern France, southern Italy, and Greece), followed by most low-altitude regions in central and eastern Europe and later by the northern Iberian Peninsula, northern France, southern England, and some higher altitude regions (Fig. 4. a-f; Supp. Video 1). For the winter chanterelle, the early days of the year show moderate probabilities of fruiting bodies occurrence in southernmost regions (e.g., Portugal and Sardinia). Predicted values then drop across Europe before increasing around mid-July in the Alps, followed by most of Northern and Eastern Europe by mid-September, and expanding to southern regions thereafter (Fig. 3 g-l; Supp. Video 2).

**Figure 4.**
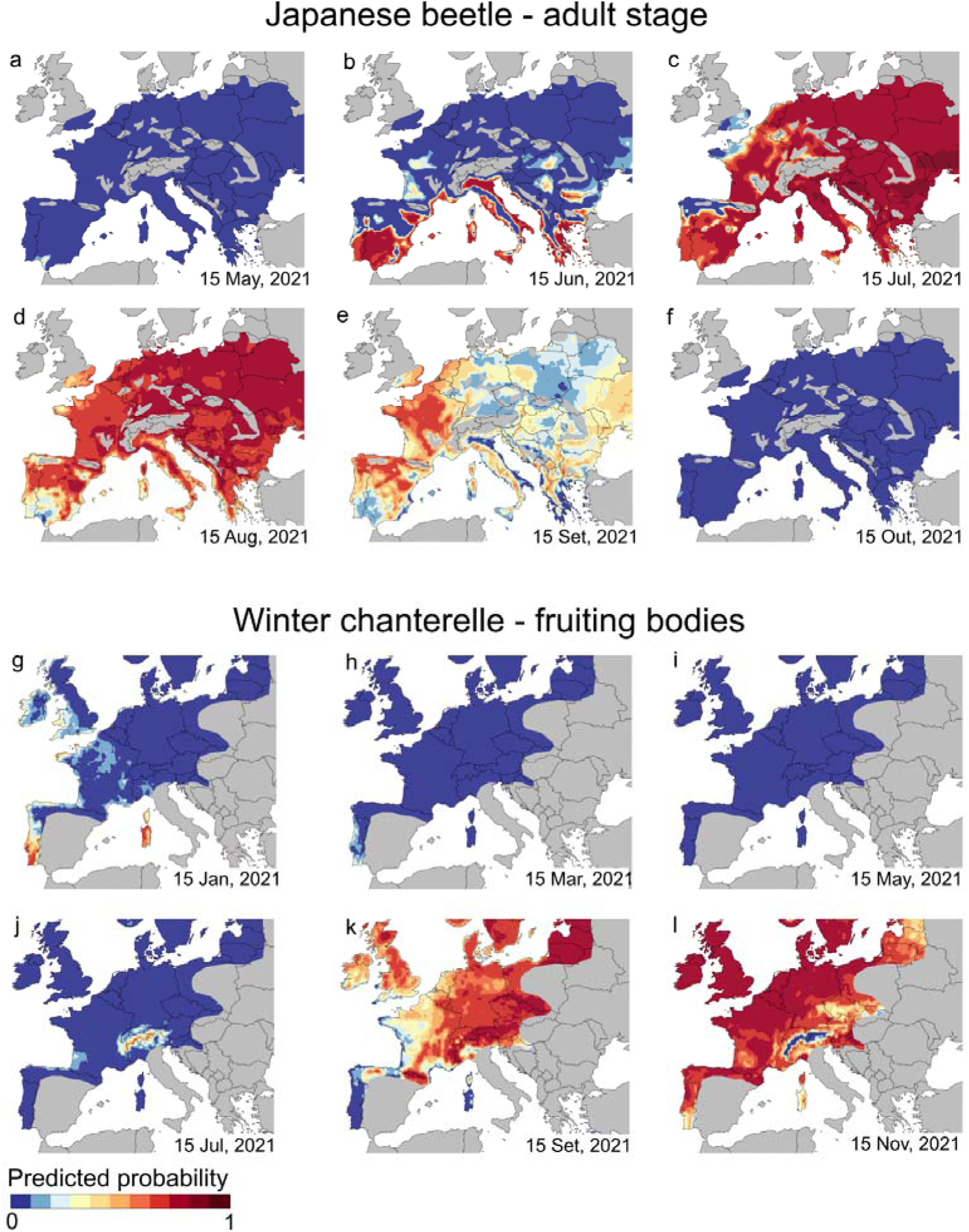
Predictions of the occurrence of adult Japanese beetles (*Popillia japonica*) (a-f) and of fruiting bodies of the winter chanterelle (*Craterellus tubaeformis*) (g-l) across Europe in selected days of 2021. Predictions are based on models trained on observation data corrected for spatial and temporal bias and AgERA5 weather data. Areas in grey are expected to be unsuitable for the Japanese beetle (a-f) or are outside the distribution range of observation records of the winter chanterelle (g-l).

## Discussion

We presented a methodological approach that allows predicting the timing of ecological events over wide geographical areas using opportunistic observation data, such as the data typically gathered from citizen science initiatives. The approach is theoretically grounded and was applied to predict the emergence period of adult Japanese beetles, an invasive species, and the availability of winter chanterelle fruiting bodies, an edible mushroom, across North America and Europe.

The approach demonstrated good predictive performance and strong agreement with observed patterns for both ecological phenomena. Based on the values of AUC – measuring the agreement between predictions and record labels for years left out of model training – the models for Japanese beetle appear more robust than those of the winter chanterelle. However, the lower AUC values achieved for the winter chanterelle may be partly attributable to its longer season of suitable conditions, which extends up to approximately 5 months, compared to the 2.5 to 3 months for the Japanese beetle. This longer season results in the generation of a higher number of temporal pseudo-absences during periods that are environmentally suitable, thereby increasing the misclassification of these records in the evaluation datasets (Philips et al., 2009). Relevantly, the predictions were still robust when made to new areas, i.e., under spatial transferability, a gold benchmark for spatial prediction in ecology (Roberts et al., 2017). This approach could be used to determine the optimal timing for surveillance efforts, particularly in the case of the Japanese beetle, a species that has not become established in most of Europe yet.

Although spatial and temporal biases in event observation data are not central to our modelling approach, they are a major source of contention in the development of predictions and estimates in temporal ecological research (Isaac and Pocock, 2015). To address these biases, we proposed and tested a set of procedures based on the patterns observed for a ‘benchmark’ taxonomic group, which is believed to represent observer bias rather than taxon-specific phenology variation. Our results show that models accounting for these biases did not differ meaningfully in their predictive performance from those that did not account for them. This is because the models estimate the probability of a set of conditions being within relative phenological niches, rather than temporal trends *per se*. In other words, the models estimate the suitability of conditions based on sampling data that does not need to be collected systematically across time and space. Instead, the representativeness of the data emerges from the joint sampling of conditions across regions and time periods. Therefore, although observational data may be biased and sparse in parts of its range, the combined use of all available observation records, representing suitable conditions, may allow for sampling most of the phenological niche.

Despite its demonstrated capability, there are several opportunities for future improvement of this approach. For instance, future work could explore general issues related to data-driven modelling, such as exploring additional predictive algorithms or different values in their parameterization. Additionally, certain design choices could be further explored and optimized, such as the number of pseudo-absences to be extracted per observation record or the procedures used to translate environmental drivers into temporally discrete predictors. For more complex phenomena, such as those with temporal dependencies or interactions between events, framework extensions may also be necessary. Specifically, recursive fitting of models, i.e., fitting the models with predictions for past periods could allow accounting for the temporal dependencies of specific phenological responses (Staggemeier et al., 2020). Similarly, predictions of interacting events could be performed and jointly modelled (Schermer et al., 2020). In addition, given the rapid pace of environmental change, it also seems essential to continuously calibrate and update these models using the most recent observation data through iterative modelling (Dietze et al., 2018). Given the emergence of novel environmental combinations in regions where the phenomena are observed, the lack of sampling of phenological responses under those settings may cause the models to fail. Therefore, continuous calibration using the most recent data is crucial to ensure that the models remain relevant and accurate over time.

## Conclusion

Our methodological approach allows obtaining informative predictions of the timing of ecological events over wide geographical areas. We demonstrated its effectiveness in predicting two distinct ecological phenomena, namely Japanese beetles and winter chanterelle mushrooms. The potential applications are vast, particularly considering the growing volumes of opportunistic observation data that are now available from various citizen science platforms. Further development and implementation of this approach are likely to make significant contributions to management-related activities such as ecological risk assessment, natural resource management, and conservation planning.

## Acknowledgments

This work was developed within the scope of project EuropaBON project, funded by European Union’s Horizon 2020 research and innovation programme under grant agreement No 101003553. C.C. also acknowledges support from Portuguese national funds provided by FCT – Fundação para a Ciência e a Tecnologia, I.P. – to the CEG/IGOT Research Unit (UIDB/00295/2020 and UIDP/00295/2020). Sergio de Miguel benefitted from a Serra-Húnter fellowship provided by Generalitat de Catalunya. M.P. was supported by national funds through FCT – Fundação para a Ciência e a Tecnologia, I.P. in the scope of Norma Transitória – DL57/2016/CP1440/CT0017.

## Author Contributions

C.C. conceptualized the study in collaboration with all authors and led the manuscript preparation with input from all authors. C.C. also performed the data analysis. A.C. and C.C. collected and visually classified the observation records.

## Data Availability

Spatial weather data used for this work are publicly available online from Copernicus Climate Data Store and National Centers for Environmental Prediction. Event observation data are available from GBIF: https://doi.org/10.15468/dl.n2agbd; https://doi.org/10.15468/dl.cc6bbn; https://doi.org/10.15468/dl.jeq9wa and https://mushroomobserver.org.

## Notes

### Competing Interest Statement

The authors have declared no competing interest.

