## Supplementary Information for "Predicting the timing of ecological phenomena across regions using citizen science data"

| **Table S1.** List of the 67 features used to characterize temporal environmental conditions for observation and temporal pseudo-absence records. In features description ‘T’ stands for the date of the observation records and the associated numbers represent preceding days. | | | | | | | | | | | | | |
| --- | --- | --- | --- | --- | --- | --- | --- | --- | --- | --- | --- | --- | --- |
| **Geography** |  |  | **Mean temperature** |  | **Minimum temperature** |  | **Maximum temperature** |  | **Precipitation** |  | **Snow depth** |  | **Wind speed** |
| Latitude |  |  | Mean T -1 to T -2 |  | Mean T -1 to T -7 |  | Mean T -1 to T -7 |  | Sum T -1 to T -2 |  | Mean T -1 to T -5 |  | Mean T -1 to T -3 |
| Longitude |  |  | Mean T -1 to T -5 |  | Mean T -8 to T -14 |  | Mean T -8 to T -14 |  | Sum T -1 to T -4 |  | Mean T -1 to T -15 |  | Mean T -1 to T -7 |
|  |  |  | Mean T -6 to T -10 |  | Mean T -15 to T -21 |  | Mean T -15 to T -21 |  | Sum T -5 to T -8 |  | Mean T -16 to T -30 |  | Mean T -8 to T -14 |
|  |  |  | Mean T -11 to T -15 |  |  |  |  |  | Sum T -9 to T -12 |  |  |  |  |
|  |  |  | Mean T -16 to T -20 |  |  |  |  |  | Sum T -13 to T -16 |  |  |  |  |
|  |  |  | Mean T -21 to T -30 |  |  |  |  |  | Sum T -1 to T -8 |  |  |  |  |
|  |  |  | Mean T -31 to T -40 |  |  |  |  |  | Sum T -9 to T -16 |  |  |  |  |
|  |  |  | Mean T -41 to T -50 |  |  |  |  |  | Sum T -17 to T -24 |  |  |  |  |
|  |  |  | Mean T -51 to T -60 |  |  |  |  |  | Sum T -25 to T -32 |  |  |  |  |
|  |  |  | Mean T0 to T -29 |  |  |  |  |  | Sum T -33 to T -47 |  |  |  |  |
|  |  |  | Mean T -30 to T -59 |  |  |  |  |  | Sum T -48 to T -62 |  |  |  |  |
|  |  |  | Mean T -60 to T -89 |  |  |  |  |  | Sum T0 to T -29 |  |  |  |  |
|  |  |  | Mean T -90 to T -119 |  |  |  |  |  | Sum T -30 to T -59 |  |  |  |  |
|  |  |  | Mean T -120 to T -149 |  |  |  |  |  | Sum T -60 to T -89 |  |  |  |  |
|  |  |  | Mean T -150 to T -179 |  |  |  |  |  | Sum T -90 to T -119 |  |  |  |  |
|  |  |  | Mean T -274 to T -364 |  |  |  |  |  | Sum T -120 to T -149 |  |  |  |  |
|  |  |  | Mean T -182 to T -273 |  |  |  |  |  | Sum T -150 to T -179 |  |  |  |  |
|  |  |  | Mean T -182 to T -364 |  |  |  |  |  | Sum T -180 to T -209 |  |  |  |  |
|  |  |  | Mean T0 to T -365 |  |  |  |  |  | Sum T -274 to T -364 |  |  |  |  |
|  |  |  | Growing degree days since 1st Julian day (baseline 0ºC) |  |  |  |  |  | Sum T -182 to T -273 |  |  |  |  |
|  |  |  | Growing degree days since 1st Julian day (baseline 7ºC) |  |  |  |  |  | Sum T -182 to T -364 |  |  |  |  |
|  |  |  | Growing degree days since 1st Julian day (baseline 15ºC) |  |  |  |  |  | Sum T0 to T -365 |  |  |  |  |
|  |  |  | Growing degree days of past 90 days (baseline 2ºC) |  |  |  |  |  |  |  |  |  |  |
|  |  |  | Growing degree days of past 60 days (baseline 2ºC) |  |  |  |  |  |  |  |  |  |  |
|  |  |  | Growing degree days of past 30 days (baseline 2ºC) |  |  |  |  |  |  |  |  |  |  |
|  |  |  | Growing degree days of past 30 to 15 days (baseline 2ºC) |  |  |  |  |  |  |  |  |  |  |
|  |  |  | Growing degree days of past 15 days (baseline 2ºC) |  |  |  |  |  |  |  |  |  |  |
|  |  |  | Growing degree days of past 7 days (baseline 2ºC) |  |  |  |  |  |  |  |  |  |  |
|  |  |  | Cold accumulation since 1st Julian day (baseline 5ºC) |  |  |  |  |  |  |  |  |  |  |
|  |  |  | Cold accumulation since 1st Julian day (baseline 10ºC) |  |  |  |  |  |  |  |  |  |  |
|  |  |  | Cold accumulation of past 30 days (baseline 5ºC) |  |  |  |  |  |  |  |  |  |  |

**Table S2.** Results of a generalized linear model with binomial error distribution, relating the presence or absence of records of the benchmark taxonomic group (*Pinus* spp.) and predictors representing calendar- and weather-related conditions for matching days.

|  | Estimate | Std. Error | *P* | Sig. |
| --- | --- | --- | --- | --- |
| (Intercept) | -9.349 | 0.128 | < 2e-16 | *** |
| Monday | -0.032 | 0.010 | 1.81E-03 | ** |
| Saturday | 0.246 | 0.010 | < 2e-16 | *** |
| Sunday | 0.177 | 0.010 | < 2e-16 | *** |
| Thursday | -0.013 | 0.010 | 1.90E-01 |  |
| Tuesday | -0.042 | 0.010 | 3.16E-05 | *** |
| Wednesday | 0.036 | 0.010 | 3.22E-04 | *** |
| August | 0.180 | 0.013 | < 2e-16 | *** |
| December | -0.658 | 0.015 | < 2e-16 | *** |
| February | -0.603 | 0.015 | < 2e-16 | *** |
| January | -0.563 | 0.015 | < 2e-16 | *** |
| July | 0.518 | 0.013 | < 2e-16 | *** |
| June | 0.187 | 0.013 | < 2e-16 | *** |
| March | -0.354 | 0.014 | < 2e-16 | *** |
| May | 0.129 | 0.012 | < 2e-16 | *** |
| November | -0.457 | 0.014 | < 2e-16 | *** |
| October | -0.168 | 0.013 | < 2e-16 | *** |
| September | -0.029 | 0.013 | 2.30E-02 | * |
| Mean temperature | 0.033 | 0.000 | < 2e-16 | *** |
| Total precipitation | -0.020 | 0.001 | < 2e-16 | *** |
| Mean wind speed | -0.013 | 0.002 | 1.61E-13 | *** |
| Significance codes: ‘***’ 0.001; ‘**’ 0.01; ‘*’ 0.05 | | | |  |

**Supplementary Figures**

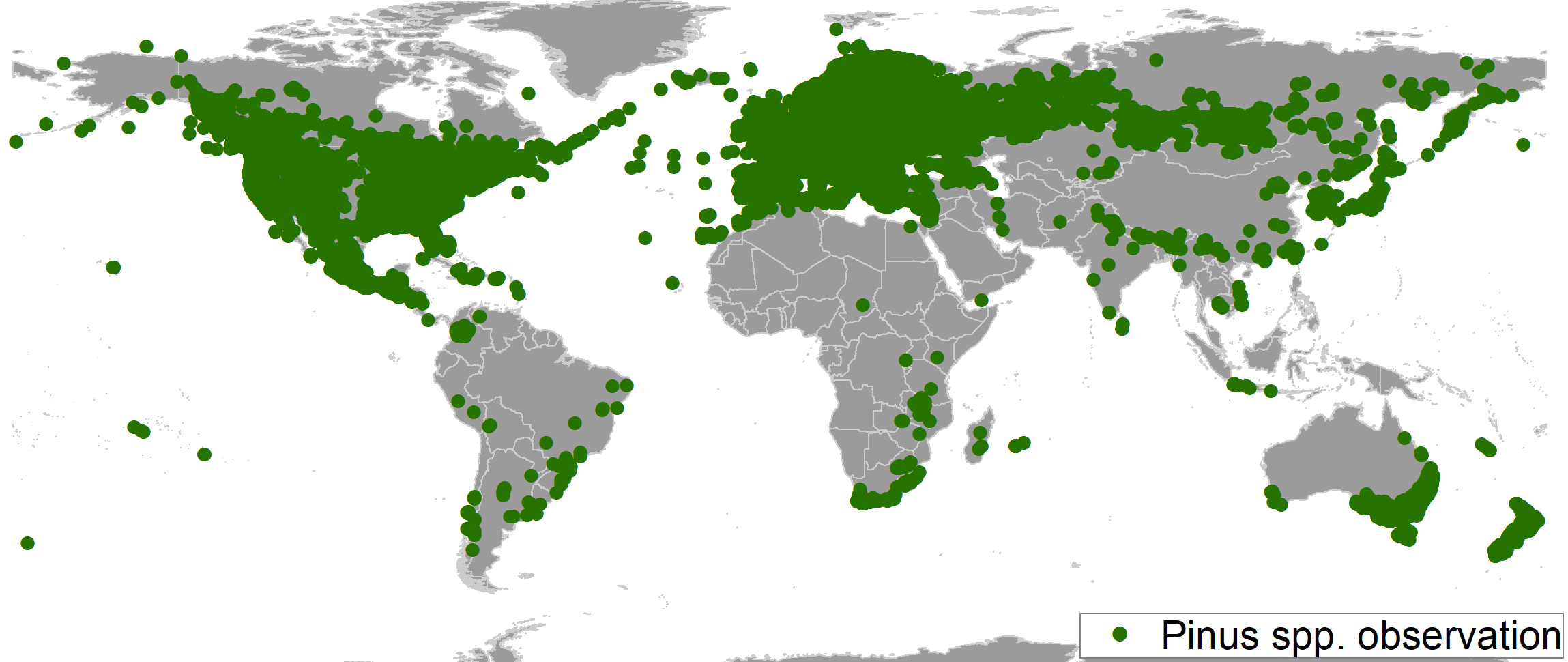

**Figure S1.** Distribution of observation records of *Pinus* spp. used to identify patterns of temporal bias in recording effort.

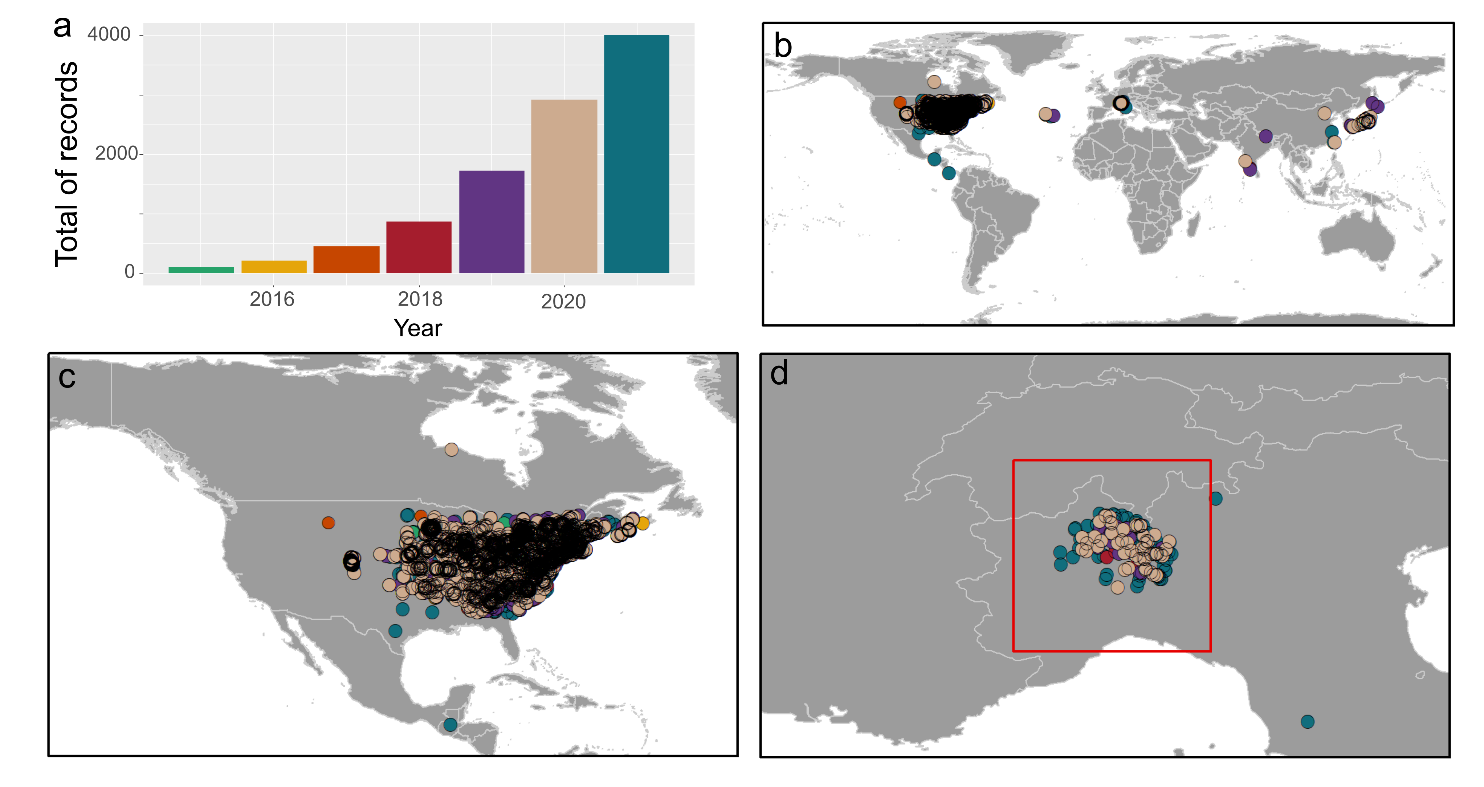

**Figure S2.** Temporal (a) and spatial (b-d) distribution of observation records of the Japanese beetle (*Popillia japonica*), after accounting for geographic overrepresentation, i.e., used for modelling. The red square in panel d delimits the observation records in Italy that were used for model validation in this region.

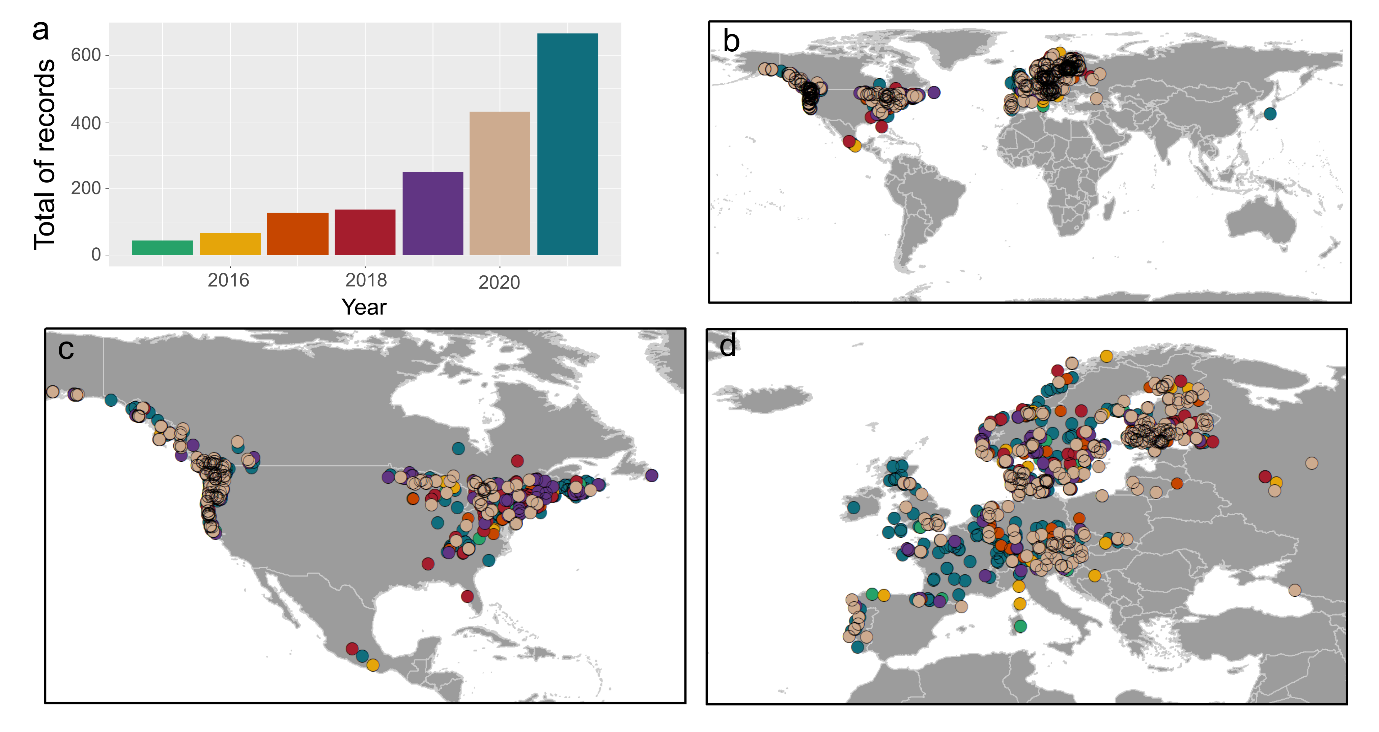

**Figure S3.** Temporal (a) and spatial (b-d) distribution of observation records of the winter chanterelle (*Craterellus tubaeformis*), after accounting for geographic overrepresentation, i.e., used for modelling.

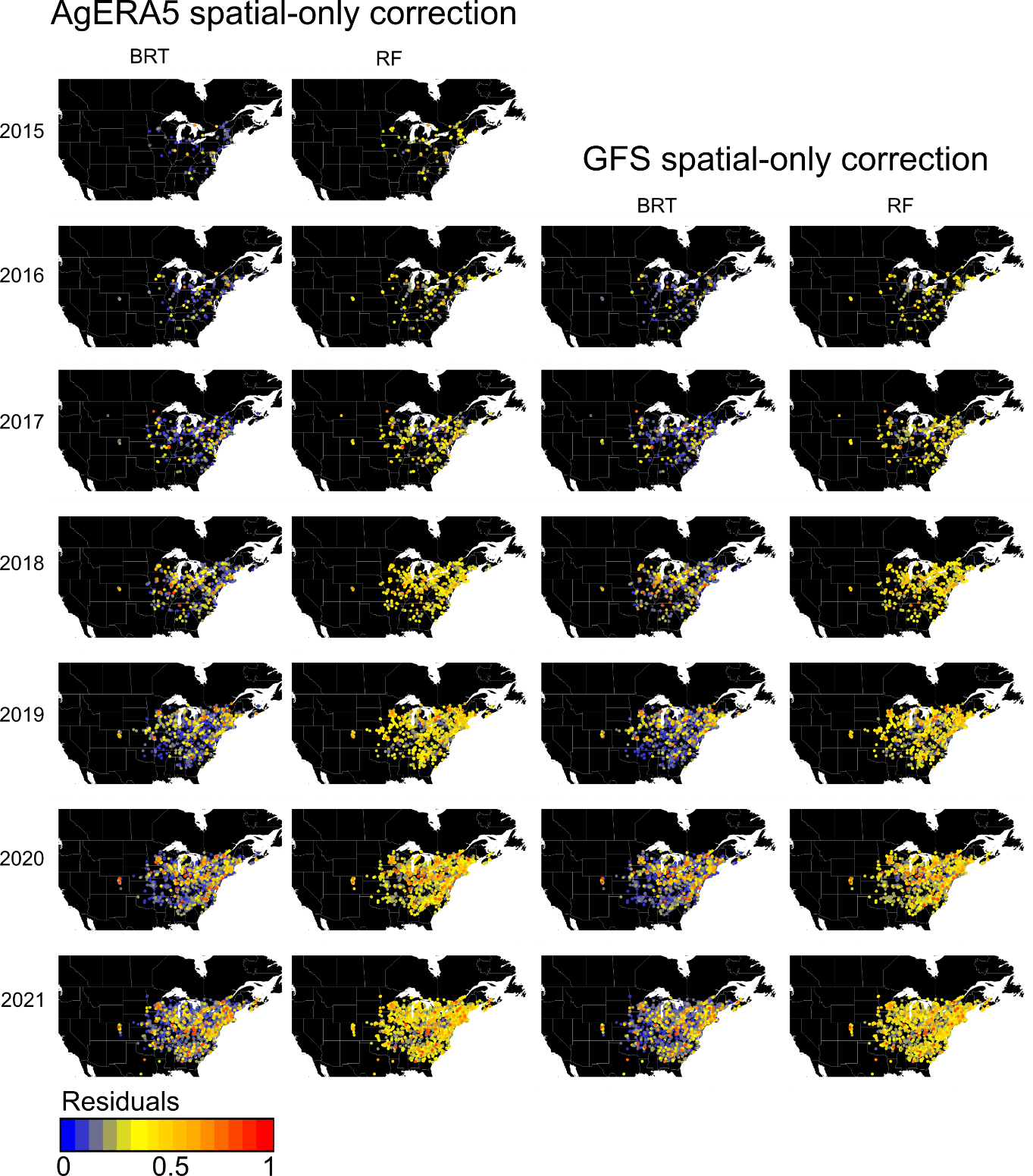

**Figure S4.** Residual values of models for the Japanese beetle (*Popillia japonica*) trained on observation data without temporal bias correction.

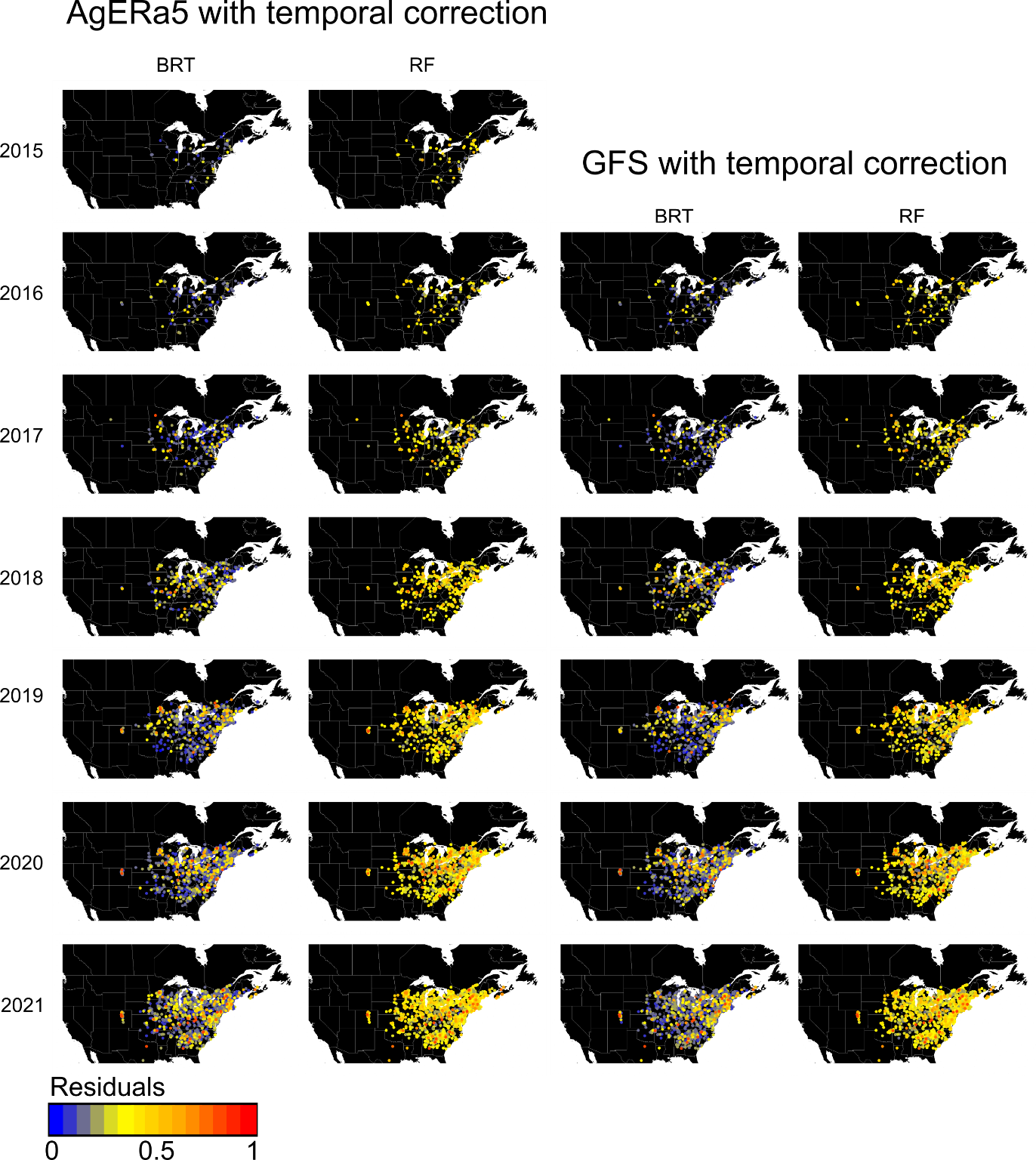

**Figure S5**. Residual values of models for the Japanese beetle (*Popillia japonica*) trained on observation data with temporal bias correction.

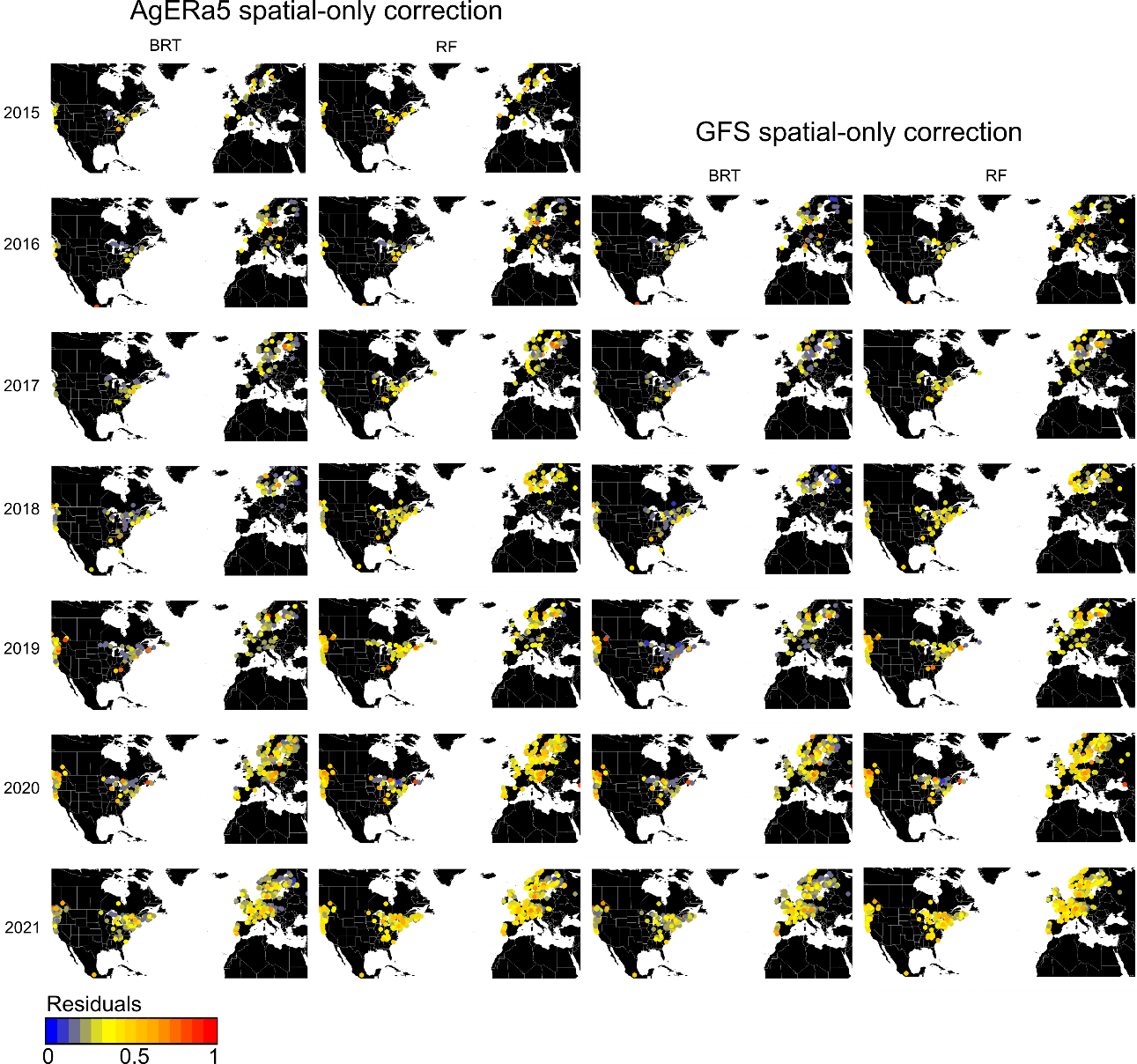

**Figure S6.** Residual values of models for the winter chanterelle (*Craterellus tubaeformis*) trained on observation data without temporal bias correction.

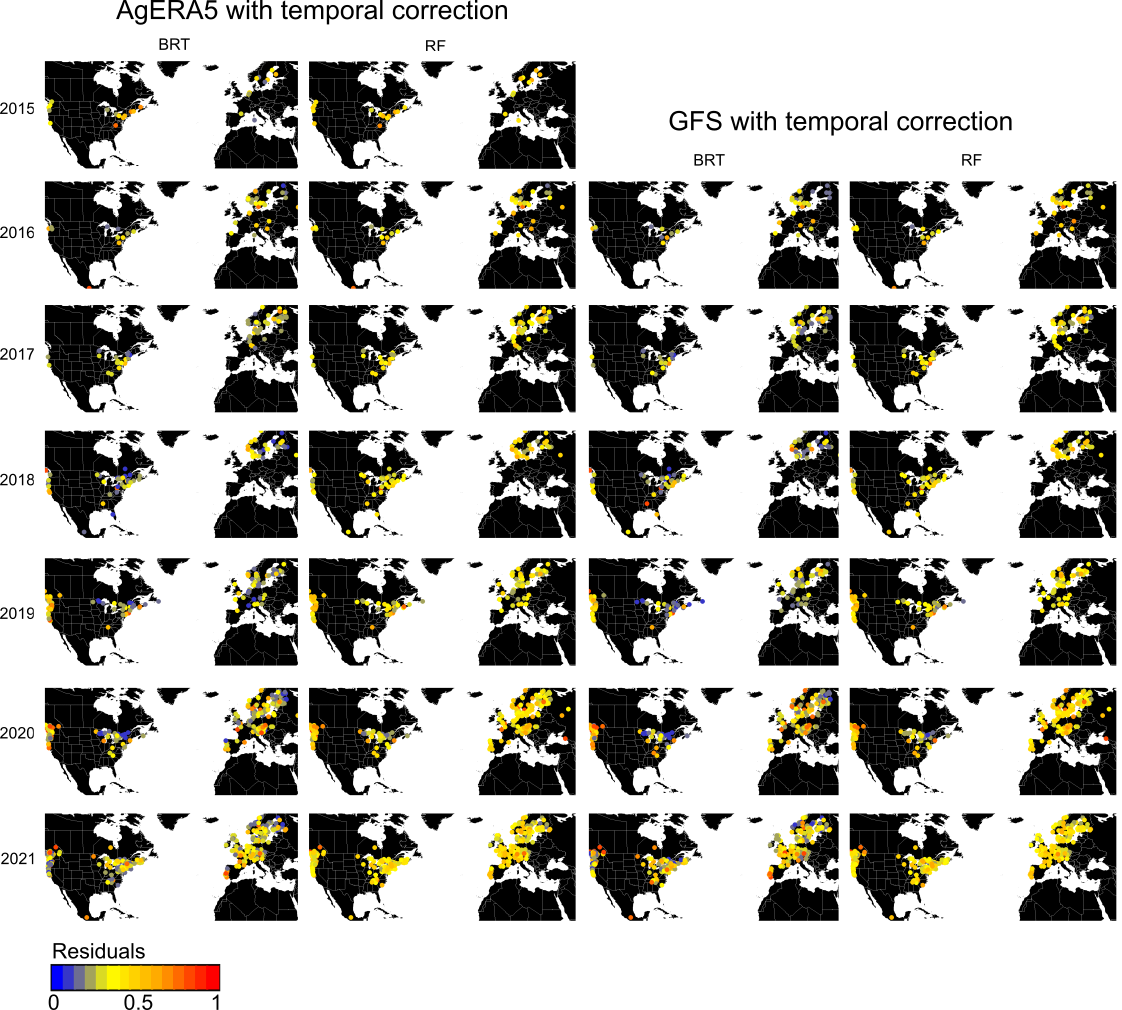

**Figure S7.** Residuals of models for the winter chanterelle (*Craterellus tubaeformis*) trained on observation data with temporal bias correction.
